# Organization of functional brain networks’ architecture during negative movie watching in late adulthood

**DOI:** 10.64898/2026.04.13.717690

**Authors:** Sara Sarebannejad, Shuer Ye, Maryam Ziaei

## Abstract

Most evidence on age-related network topology derives from resting-state paradigms, leaving unclear how aging alters brain organization during naturalistic processing and whether graph-theoretical metrics relate to emotional and cognitive functioning in ecologically valid contexts. We analyzed movie-fMRI and behavioral data from 72 younger and 68 older adults, examining global (small-worldness, clustering coefficient, characteristic path length), network (participation coefficient), and nodal (degree centrality, betweenness centrality, nodal efficiency) properties. Regression models were used to test associations between nodal measures and both the Emotional Resilience Index (ERI) and the Cognitive Function Index (CFI), while mediation analyses were conducted to test whether nodal measures mediate the relationship between age and ERI. Older adults exhibited increased characteristic path length and clustering coefficient, indicating reduced global integration and greater local segregation. Although small-world organization was preserved in two groups, there was less pronounced small-world architecture in older adults compared to younger adults, suggesting a shift toward more regularized, locally clustered networks and reduced long-range connections during dynamic stimuli. Participation coefficient values were higher in the somatomotor, frontoparietal, and default mode networks, and lower in the subcortical network, among older adults reflecting greater between-network integration in cortical networks but diminished subcortical coordination in aging. Five key nodes, two thalamic regions, hippocampus, and two insular regions, showed reduced centrality and efficiency in older adults during the negative movie, indicating weakened dominance of subcortical hubs under emotional salience condition. Right thalamic nodal properties were negatively associated with ERI and CFI and served as mediators in the relationship between age and emotional resilience. These findings suggest that reduced thalamic hub centrality may reflect adaptive recalibration of salience emotional processing, linking network reorganization to improved emotional resilience in aging.

**Key points:**

- Older adults showed higher path length and clustering, suggesting reduced integration.
- Reduced small-worldness reflects weaker balance of segregation and integration with age.
- Older adults showed higher cortical but lower subcortical participation coefficients.
- Key nodes showed reduced centrality during negative stimuli, indicating weaker hubs.
- Right thalamus changes linked to resilience, mediating age-emotion relationships.

## Introduction

Aging has been consistently associated with declines across multiple cognitive domains, including attention (Lezak et al., 2012), working memory (Kennedy et al., 2015), episodic memory, and processing speed (Deary et al., 2009). These age-related cognitive changes are closely paralleled by alterations in brain structure and function, such as reductions in gray matter volume (Raz et al., 2005), and reduced neural specificity or differentiation (Hedden & Gabrieli, 2004; Park et al., 2004; Koen & Rugg, 2019). Despite these declines, the aging brain also demonstrates adaptive responses. Compensatory neural mechanisms manifest as increased activity in the bilateral and anterior brain regions to help maintain performance, as described by prominent neurocognitive aging models, including the Posterior-Anterior Shift in Aging (PASA; Davis et al., 2008; Dolcos et al., 2002), the Hemispheric Asymmetry Reduction in Older Adults (HAROLD; Cabeza, 2002; Cabeza et al., 2002), and the Compensation-Related Utilization of Neural Circuits Hypothesis (CRUNCH; Reuter-Lorenz & Cappell, 2008). In contrast to the well-documented declines in cognition, emotional functioning appears to remain relatively preserved, or even improve, with age (Charles, 2010; Charles & Carstensen, 2010). Older adults often exhibit a positivity bias and tend to invest more in emotionally meaningful goals and relationships, patterns described by the Socioemotional Selectivity Theory (SST; Carstensen et al., 1999; Mather & Carstensen, 2005; Reed & Carstensen, 2012). Neuroimaging evidence further supports complex interplay between aging, neural adaptation and emotional stability. For instance, functional connectivity between the right thalamus or right hippocampus and anterior insula, mediate the relationship between age and positive emotional reactivity (Niu et al., 2024), highlighting the importance of subcortical-limbic regions in the relationship between emotional regulation and aging.

Age-related functional connectivity alterations reflect large-scale connectomes reorganization, rather than isolated regional effects (Zuo et al., 2012; Geerligs et al., 2015). Graph theory offers a comprehensive framework by modeling the brain as a network of nodes (brain regions) and edges (connections between regions), enabling quantification of age-related reconfiguration at global, network, and nodal levels (Rubinov & Sporns, 2010). At the global level, clustering coefficient and characteristic path length indicate local segregation and global integration, respectively, with their balance captured by small-world architecture (Watts & Strogatz, 1998; Sporns, 2013). Although small-worldness is typically preserved in healthy brains, but reduction in small-world organization is identified in older adults (Onoda & Yamaguchi, 2013; Xu et al., 2015; Q. Wang et al., 2024). Some studies reporting increased clustering coefficients and path lengths in aging (Sala-Llonch et al., 2014; Xu et al., 2015), whereas others show divergent patterns (Hugenschmidt et al., 2014; Bagarinao et al., 2019; Yu et al., 2025). At the network level, participation coefficient quantifies inter-module connectivity (Guimerà & Amaral, 2005). Increased between-network integration (Dennis & Thompson, 2014; Deery et al., 2023), and greater inter-module connectivity across cortical networks occur in aging (Chan et al., 2014; Chong et al., 2019; Madden et al., 2024). At the nodal level, centrality and efficiency metrics identify critical hubs in the brain’s overall organization (Achard & Bullmore, 2007; Rubinov & Sporns, 2010), to facilitate communication between disparate regions (van den Heuvel & Sporns, 2013). Aging has been associated with reduced nodal importance in regions such as the bilateral amygdala, left thalamus, left insula and right hippocampus and right parahippocampus (Achard & Bullmore, 2007), alongside increased centrality in the frontal areas (Lee et al., 2016; Behfar et al., 2020), suggesting that the reduction in the significance of subcortical-limbic nodes, anterior redistribution and hub reorganization across the lifespan. Collectively, these prior findings indicate heterogeneous and sometimes inconsistent alterations in topological findings across the lifespan, which could also be related to different brain parcellations schemes, sparsity thresholds, and brain atlases used. In addition, while some studies link graph measures to cognitive performance in late adulthood (Lee et al., 2016; Baeuchl et al., 2019; Chong et al., 2019; Behfar et al., 2020; Yao et al., 2025), it remains unclear whether age-related differences in all three different levels of graph properties are associated with age-related cognitive or emotional well-being functioning.

In recent years, naturalistic paradigms, such as movie-functional magnetic resonance imaging (movie-fMRI), are increasingly gaining prominence due to their ability to capture ecologically valid patterns of brain activity (Hasson et al., 2010; Eickhoff et al., 2020). Compared with traditional task-based paradigms, movie-fMRI enables the acquisition of high-quality neural data by evoking naturalistic cognitive and emotional responses (Hasson et al., 2004; Cantlon & Li, 2013) while reducing confounding effects associated with task repetition and explicit performance demands (Vanderwal et al., 2019). In movie-fMRI paradigms, participants view films passively, a design that promotes sustained engagement (Spiers & Maguire, 2007), reduces head motion (Dosenbach et al., 2017), and minimizes the risk of sleep during scans (Horovitz et al., 2008), factors particularly critical in aging research, where compliance and fatigue can bias resting-state estimates. Importantly, movie-fMRI captures ongoing fluctuations in attention, affect, and salience processing, thus, engaging large-scale integration mechanisms that may not be fully expressed during rest (Meer et al., 2020; Jääskeläinen et al., 2021; Song et al., 2023; Ke et al., 2025). Consistent with this, connectivity patterns derived from movie-fMRI predict chronological age more accurately than resting state fMRI (rs-fMRI), suggesting greater sensitivity to age-related network reconfiguration (Bi et al., 2024). Furthermore, functional connectivity during movie watching yields better predictions than rs-fMRI in trait behaviors, in both the cognition and emotion domains (Finn & Bandettini, 2021). Emotional stimuli further modulate connectivity in subcortical–limbic regions, including the bilateral amygdala, thalamus, putamen, and hippocampus (Kinreich et al., 2011), and can identify individuals with emotional vulnerability (Ye et al., 2026). Negatively-valanced stimuli can reveal deficits in brain organization that are underestimated under low-demand or neutral conditions (Kirk et al., 2023; Wang et al., 2023). Despite these advantages, graph-theoretical analyses have rarely been applied to movie-fMRI in aging populations. The present study, therefore, leverages movie-fMRI to examine age-related differences in brain network topology under realistic conditions. We hypothesize that emotionally engaging stimuli will reveal behaviorally relevant reconfiguration of large-scale networks in older adults, particularly within subcortical and regulatory hubs, beyond what is detectable in younger adults or resting paradigms reported in previous studies.

The present study aims to investigate age-related differences in functional brain network topology at the global, network, and nodal levels, using ecologically valid movie-fMRI with two emotionally valanced conditions. The current study extends this framework to examine association between topological measures and emotional functioning in older adults, an area that remains largely unexplored. First, we hypothesized that older adults would show systematic reconfiguration of large-scale brain topology during naturalistic processing, characterized by decreased small-worldness (reflecting reduced balance of local segregation and global integration), greater participation coefficient values of the cortical networks (reflecting greater inter-module connectivity of the cortical networks), and reduced dominance of subcortical-limbic areas, as hub regions, and increased centrality of prefrontal areas. Such pattern is expected to reflect age-related weakening of long-range integration reflected in characteristic path length, stronger inter-connectivity across cortical brain networks, and redistribution of hub roles. Second, we expected that network topology would be differentially associated with emotional resilience and cognitive functioning. Specifically, we predicted that greater balance of local specialization and global integration (i.e., greater small-worldness), reduced inter-module integration, and preserved hub nodes in key regulatory regions, such as prefrontal areas, would be associated with higher cognitive function, while less dominance in subcortical regions are linked to higher emotional resilience, reflecting adaptive salience processing in aging. Third, we hypothesized that nodal properties of subcortical hubs, such as the thalamus, hippocampus and insula, would mediate the relationship between age and emotional resilience, while the centrality of frontal areas would be accounted for the association between age and executive functioning. It is predicted that age-related nodal reorganization contributes to preserved emotional functioning in late adulthood.

## Materials and methods

### Study 1: Mini literature review on aging and topological measurements

To clarify the magnitude of age effects in graph theoretical measures, we first conducted a literature review to identify all relevant studies for this study and to identify what graph measures to be used for our analysis. The search was performed on PubMed in April 2025, supplemented by reference tracking from previous systematic reviews and meta-analyses. The search terms included the following: (“aging” OR “ageing” OR “aged” OR “older” OR “elder” OR “elderly” OR “lifespan”) AND (“fMRI” OR “functional magnetic resonance imaging” OR “functional MRI” OR “resting state” OR “task-related” OR “movie-fMRI” OR “movie-watching”) AND (“graph theory” OR “graph analysis”). A total of 1,969 records were screened, and studies were included based on the following criteria: (1) inclusion of subjects over 18 years old; (2) absence of any neurological or psychiatric disorders; (3) investigation of brain functional topology based on the predefined graph measures (Rubinov & Sporns, 2010) (4) Assessment of age effect or the comparison between older and younger adult groups; and (5) inclusion of older subjects aged 65 years or above. Systematic reviews, meta-analyses, case reports, editorial letters, and methodological studies were excluded. Following a thorough review, 23 publications met the inclusion criteria. A summary of the findings is presented in Table S1.

### Study 2: Age-related functional reorganization during movie-fMRI

#### Participants

The data used in this study were collected as part of the larger Trondheim Aging Brain Study (TABS), which has been described in detail elsewhere (Dave et al., 2025; Ye et al., 2025). Initially, a total of 80 healthy older adults and 80 younger adults were recruited from the local community in Trondheim through paper flyers and online advertisements. Inclusion criteria comprised right-handedness, no history of neurological disease (e.g., epilepsy, stroke, or brain injury), no diagnosis of severe psychiatric disorders (e.g., depression, anxiety, schizophrenia, or autism spectrum disorder) within the past five years, and no current use of medication affecting mood or neurological function. Following data quality assessment, twenty participants were excluded due to excessive head motion (average framewise displacement > 0.3 mm), failure to complete the experiment, or technical issues during data acquisition. This resulted in a final sample of 68 healthy older adults (36 females; mean age = 70.90 ± 3.83 years; age range: 65–82 years) and 72 younger adults (34 females; mean age = 25.75 ± 4.25 years; age range: 19–36 years) included in the analyses. This study was conducted in accordance with The Declaration of Helsinki, and ethical approval was granted by the Norwegian Regional Committee for Medical and Health Research Ethics (Midt REC). Written informed consent was obtained from all participants, and each participant received a gift card as compensation upon completion of the experiment.

#### Movie clips and procedure

A pilot behavioral study was conducted to identify and select suitable movie clips for the imaging experiment. Four candidate films, drawn from previous studies and an online repository (Taylor et al., 2017), were evaluated by twenty-four participants (12 females, mean age =25.5 ± 3.96 yrs). Participants rated the clips on emotional valence, arousal, emotion category, and continuous emotional intensity throughout the viewing period. Based on the pilot study results, two clips were selected for use in the subsequent imaging session. The neutral movie clip, “Pottery” (480 seconds in duration), depicts two women making pottery and contains relatively low emotional content, whereas the negative clip, “Curve” (494 seconds in duration), portrays a woman struggling to avoid falling into a dam, and is characterized by high salient, stress-inducing emotional content (details of the content characterization and pilot results are reported in Ye et al., (2025). To minimize potential confounds across clips, several key features were controlled: (1) no more than one human character appears on screen at any given time; (2) all characters are female; (3) the clips contain no dialogue, written text, or subtitles; and (4) both clips are presented in color. Accordingly, the primary analyses focused on the negative movie condition, which was validated as emotionally evocative, while parallel analyses of the neutral conditions are reported in the Supplementary Materials.

The experiment consisted of two sessions: a brain imaging session and a behavioral session (Figure 1). Initially, all participants underwent an MRI scan, which included both structure and functional and imaging protocols. Prior to scanning, participants were provided with verbal and written instructions regarding the experimental procedure. During the scanning procedure, participants first underwent an 8-minute anatomical scan followed by a movie-watching session, which included first the neutral, followed by the negative movie clip. Participants verbally rated their emotional valence (from very unpleasant to very pleasant) and arousal levels (from very calm to very excited), using a 9-point Self-Assessment Manikin (SAM) scale (Bradley & Lang, 1994), before and after each movie presentation. To eliminate potential confounding factors, stimuli variability was systematically controlled using fixed movie sequences with consistent contextual content (Chen et al., 2020; Baek et al., 2022). Foam padding was used to minimize head motion, and participants were provided with MRI-compatible earphones (BOLDfonic, Cambridge Research Systems Ltd) with adjusted sound for each individual prior to watching the movies. They viewed the movie clips through a mirror mounted on the head coil. Within two to four days following the brain imaging session, they completed a behavioral session lasting approximately two hours, during which they filled out self-reported questionnaires and performed computerized tasks.

**Figure 1.**
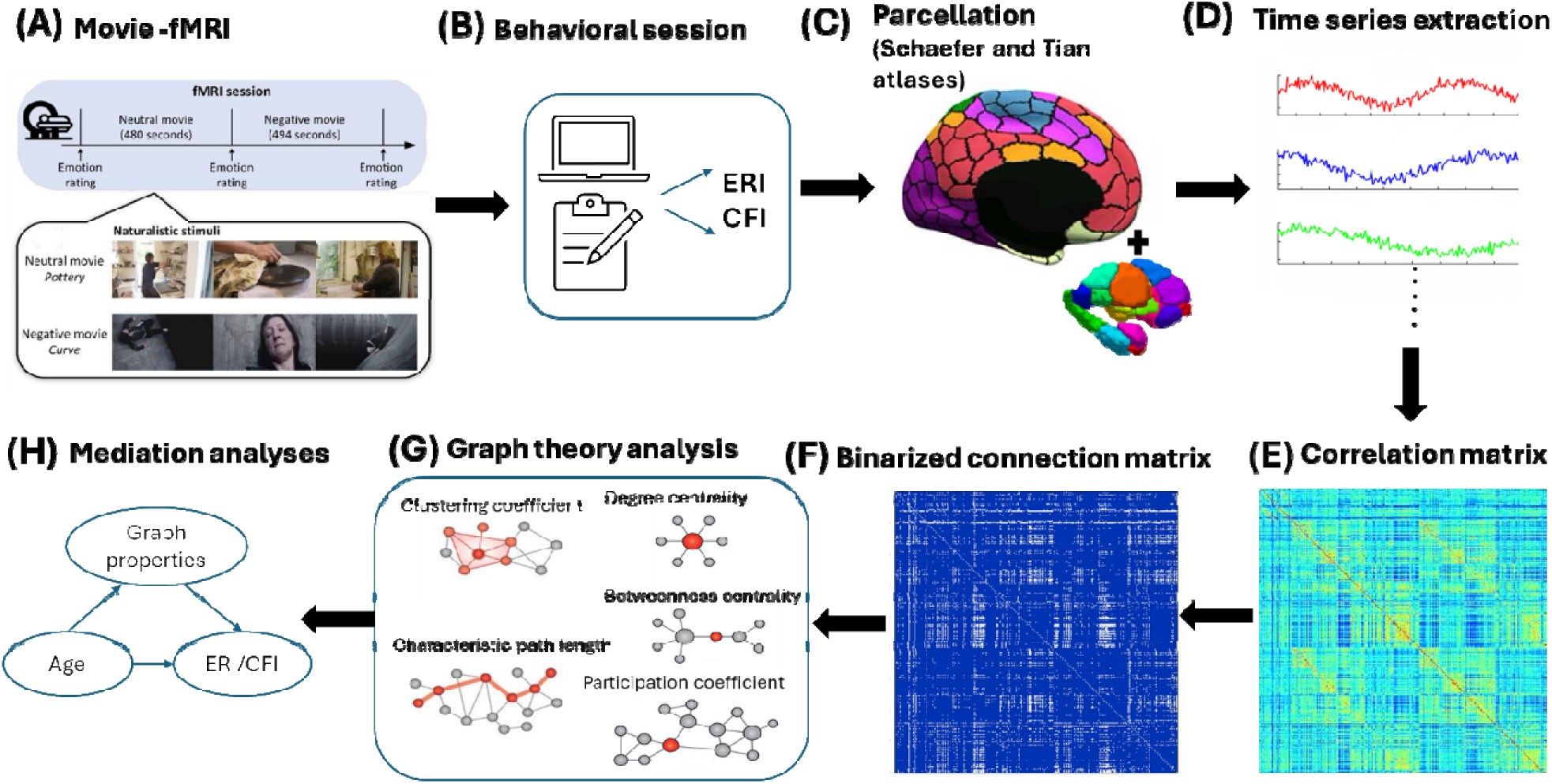
Schematic overview of analyses pipeline. After participants viewed passively neutral and negative movie in a 7 T MRI scanner (Ye et al., 2025), (A) whole-brain BOLD signals were acquired from all participants while watching neutral and negative movies, followed by (B) filling out self-reported questionnaires and performing computerized tasks (measures of the Emotional Resilience Index (ERI) and the Cognitive Function Index (CFI)). (C) Parcellation into 454 distinct cortical and subcortical brain regions using (Schaefer et al., 2018) and (7T-S3, Tian et al., 2020) for each subject. (D) Time series of voxels within the given region were extracted, (E) subsequently, average time courses of all pairs were calculated by Pearson correlation to construct 454 × 454 correlation matrices per subject. (F) The correlation matrices were binarized and thresholded to create adjacency matrices, and (G) graph theory analysis were applied to compute the global, network and nodal measures (the graphic illustration of graph properties were adopted from (Farahani et al., 2019), and (H) mediation analyses were then conducted to investigate whether the association between age and ERI/CFI are mediated by graph properties. ERI= emotional resilience index; CFI= Cognitive Function Index.

#### Neuropsychological assessment

Following the scanning session, participants completed a series of self-reported questionnaires and computer-based tasks to evaluate psychological well-being and cognitive functioning. Psychological well-being was assessed using standardized instruments including the Depression, Anxiety, and Stress Scale (DASS-21; Lovibond & Lovibond, 1995), the Difficulties in Emotion Regulation Scale (DERS; Gratz & Roemer, 2004), the General Health Questionnaire (GHQ-28; Goldberg & Hillier, 1979), the Hospital Anxiety and Depression Scale (HADS; Zigmond & Snaith, 1983), the Connor-Davidson Resilience Scale (CD-RISC; Connor & Davidson, 2003), the Intolerance of Uncertainty Scale (IUS; Freeston et al., 1994; Buhr & Dugas, 2002), the Perceived Stress Scale (PSS; Cohen et al., 1983), the State-Trait Anxiety Inventory (STAI; Spielberger, 1983). Executive control was evaluated through the Stroop Test (Jensen & Rohwer, 1966), the Trail Making Test (Parts A and B; Reitan, 2003), and the COWAT Phonemic and Semantic Verbal Fluency tasks (Newcombe, 1969), and forward and backward digit span tests. Theory of mind and empathy were assessed using the Reading the Mind in the Eyes Task (RMET; Baron Cohen et al., 2001) and the Interpersonal Reactivity Index (IRI; Davis, 1980), respectively. General intelligence was measured via the shortened version of Raven’s Progressive Matrices (Raven, 1940). Table 1 includes details on descriptives of background measures.

**Table 1:**
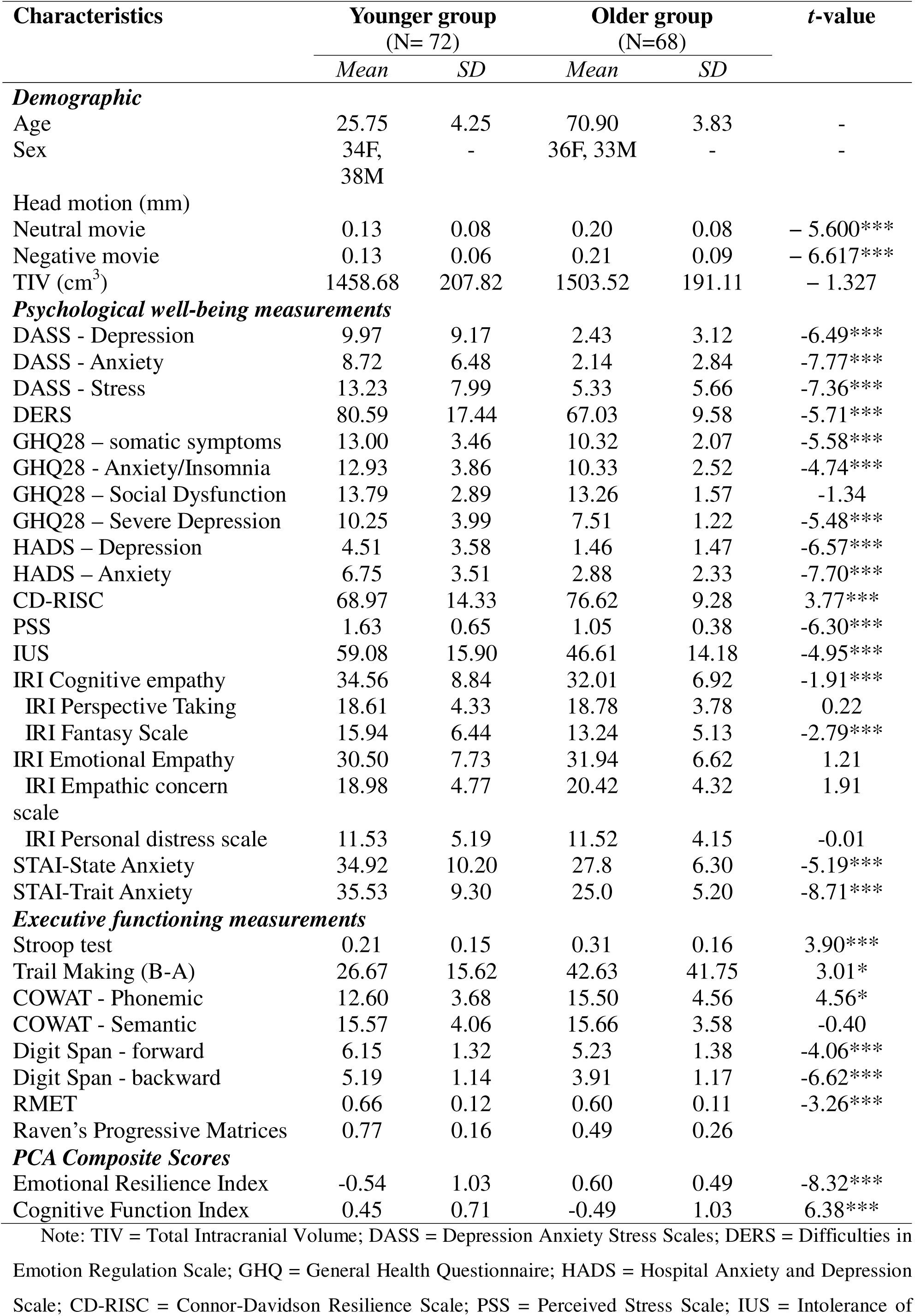
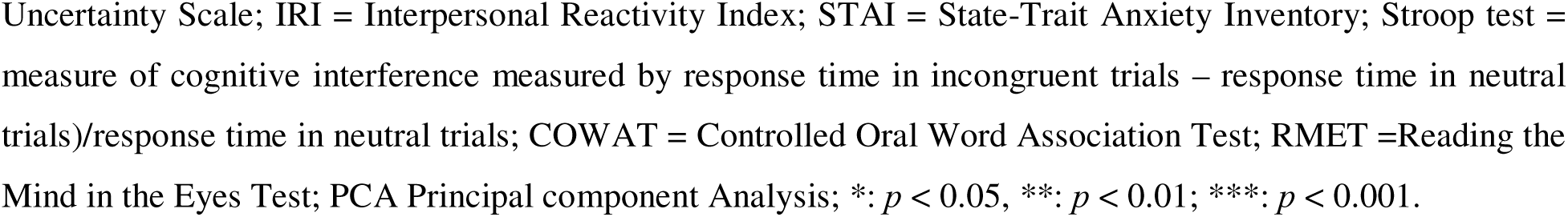
Demographic characteristics, descriptive measurements of psychological well-being questionnaires and executive functioning tasks in younger and older participants.

As multiple measures were collected, Principal Component Analysis (PCA) was performed to reduce the dimensionality of emotional and cognitive measures as in our previous work (Dave et al., 2025). The Emotional Resilience Index (ERI) was defined to capture emotional adaptability and well-being, while the Cognitive Function Index (CFI) served as a measure of cognitive performance. These indices were validated using Kaiser-Meyer-Olkin (KMO) measure of sampling adequacy and Bartlett’s test, confirming strong correlations among the variables (Masullo et al., 2021). Components with eigenvalues above 1 were considered significant: ERI comprised two components explaining 63.94% of the total variance, with the first component explaining 57.05% of the variance, while CFI consisted of three components explaining, 72.69% of the total variance, with the first and second components accounting for 32.79% and 20.06%, respectively. The ERI reflects an individual’s capacity to regulate emotional responses and adapt to uncertainty, whereas the CFI represents verbal ability, working memory, and inhibitory control. Higher index scores indicate enhanced emotional resilience and cognitive functioning.

#### Imaging acquisition and preprocessing

All imaging data were acquired using a 7T MRI Siemens MAGNETOM Terra scanner with a 32-channel head coil at the Norwegian 7T MR Center, located at St. Olav’s Hospital/Norwegian University of Science and Technology. Anatomical T1-weighted high-resolution images were obtained using the Magnetization Prepared Rapid Gradient Echo (MP2RAGE) protocol (224 sagittal slices; slice thickness: 0.75 mm, echo time: 1.99 ms, repetition time: 4300 ms, inversion times: 840 ms and 2370 ms, voxel size: 0.8 × 0.8 × 0.8 mm, flip angle: 5°/6°). Functional images were acquired using a multi-band accelerated echo-planar imaging sequence (92 interleaved slices; multi-band acceleration factor: 8, echo time: 19 ms, repetition time: 2000 ms, matrix size: 160 × 160 mm, voxel size: 1.25 × 1.25 × 1.25 mm, field of view: 200 mm, slice thickness: 1.25 mm, flip angle: 80°), producing 243 volumes for the neutral movie and 250 volumes for the negative movie.

Similar to our previous work (Ye et al., 2025), the fMRI images were preprocessed using fMRIPrep version 22.0.2 (Esteban et al., 2019) and XCP-D version 0.3.053 for post-preprocessing (Mehta et al., 2024). After removing the first three dummy scans, the preprocessing steps included slice timing correction, motion correction, co-registration of structural and functional images, and normalization to MNI space. Post processing steps included band-pass filtering between 0.009–0.08 Hz, removal of nuisance regressors and spatial smoothing with 4 mm Gaussian kernel.

#### Network construction and analysis

The whole-brain functional connectivity matrices were generated using 454 distinct areas according to Schaefer 400-parcel cortical atlas (Schaefer et al., 2018) and Tian 54-parcel subcortical atlas (7T-S3, Tian et al., 2020) for each subject. The parcels were assigned to seven functionally predefined cortical networks: VIS, SMN, dorsal attention (DAN), ventral attention (VAN), limbic network (LIM), FPN, and DMN networks (Yeo et al., 2011), in addition to one subcortical (SUB) network. Following obtaining time series of voxels within the given region, Pearson correlation was further employed on the average time courses of all pairs to construct 454 × 454 functional connectivity matrices per subject. The correlation coefficient was followed by applying Fisher’s *r-*to*-z* transformation to improve data normality, resulting in two functional connectivity matrices for both negative and neutral conditions for each subject.

To perform the graph theory analysis, the correlation matrices were binarized to facilitate interpretation (Kaiser, 2011). Adjacency matrices were thresholded based on the density of connections, thereby increasing the contrast between strong and weak connections. The objective was to set a specific sparsity level that would generate an optimal network for comparison across subjects, while excluding spurious correlations. Given that sparsity level of 0.25 is the most suitable threshold to maintain the relationship between the number of nodes and the density of edges (Laurienti et al., 2011), a sparsity threshold of 0.25 was therefore chosen. This threshold yielded networks with the greatest similarity to the expected density based on the number of nodes. Accordingly, 25% of the strongest connections were assigned a value of 1, and the remaining connections were set to 0. Due to the unclear biological basis of negative correlations (Fox et al., 2009), only positive correlations were considered, and all negative connections were set to zero. The variation in global properties was examined across additional sparsity thresholds of 0.2 and 0.3, while network and nodal properties were analyzed at the sparsity value of 0.25. Graph theoretical analyses were performed using GRETNA toolbox (J. Wang et al., 2015) in MATLAB (version R2024a).

### Graph Theoretical analyses

#### Global properties

We systematically analyzed the global properties of functional networks using three graph measures: clustering coefficient, path length, and small-worldness. Clustering coefficient quantifies the extent to which an individual node can connect to its neighboring nodes and form a triangle (Rubinov & Sporns, 2010). It indicates the prevalence of clustered connectivity around each node, serving to localize information processing and implying functional specialization (Watts & Strogatz, 1998). Characteristic path length defines the minimum number of edges required to link any two nodes in the network (Rubinov & Sporns, 2010). Path length is described as the number of steps along the route between two nodes, highlighting shorter characteristic path length represents higher global communication efficiency (Watts & Strogatz, 1998).

Small-world architecture lies between regular and random networks, characterized by a higher clustering coefficient than random networks and a shorter characteristic path length than regular lattices. To interpret clustering coefficient and path length meaningfully, they must be normalized using the same values from 100 random networks with equal numbers of nodes and degree distributions, but different wiring configurations. A high small-worldness value indicates both higher local segregation and efficient global integration of the network (Sporns & Honey, 2007).

#### Network properties

Subsequently, network properties were examined by participation coefficient after partitioning the graph into modules, this measure evaluates how nodes in each module distribute connections across other modules (Guimerà & Amaral, 2005). It characterizes whether the links of a node are localized within its own module or spread across different ones (Rubinov & Sporns, 2010), indicating between-network integration.

#### Nodal properties

Lastly, we computed nodal topological properties by three measures: degree centrality, betweenness centrality, and nodal efficiency. Degree centrality captures the degree of a node, defined as the total number of direct links to other nodes (Rubinov & Sporns, 2010), and it emphasizes a node’s centrality in information flow. Betweenness centrality assesses how often a node appears on the shortest path between any two nodes (Freeman, 1978), and it identifies key nodes that act as bridges, indicating substantial control over information flow. Nodal efficiency quantifies how efficiently a node communicates with all others, using the average inverse of shortest path lengths (Latora & Marchiori, 2001). Nodes with higher nodal efficiency are more effective at distributing information quickly. The details of relative equations for all indices used in the study can be found in Supplementary Materials.

#### Statistical analyses

To assess whether there were age-related alterations in topological levels, two-sample *t*-tests were performed to examine between-group differences in global, network and nodal graph measures. Significance values were further adjusted using the false discovery rate (FDR) correction. We further computed separate linear regression analyses to determine the effects of each graph theoretical measure on ERI/CFI, with age, sex, head motion, and total intracranial volume (TIV) included as covariates. After the brain extraction, the TIV of each participant was estimated from the T1 image using the Computational Anatomy Toolbox (CAT12) (Gaser et al., 2024). Statistical analyses were performed using SPSS (version 30.0.0).

To further explore whether the effect of age on ERI/CFI was influenced by graph theoretical measures, mediation analyses were conducted using the PROCESS macro (Hayes, 2012). The mediation models were built under two premises; first the graph measure showed age differences, and second the relevant measure was associated with behavioral performance. In each model, age was set as the independent variable, relevant graph properties were included as the mediator variables, and ERI/CFI served as the dependent variable, while controlling for covariates (Sex, head motion, and TIV). A total of three mediation models were conducted to investigate the relationship between an independent and a dependent variable while accounting for the influence of mediators. These paths included: path a (predictor to mediator), and path b (mediator to outcome), controlling for the covariates. Path c (the total effect of the predictor on the outcome) comprised two components: the indirect effect (a × b path), which refers to the impact of topological properties on the effect of age on ERI or CFI; and the direct effect (c’ path), which excludes the mediator’s influence and defines the sole effect of age on ERI or CFI. Path effects were interpreted as significant if bias-corrected 95% confidence intervals did not include zero, based on 5000 bootstrap resamples (Hayes, 2009). If the indirect effect was significant and the direct effect was not, full mediation was inferred. If both the direct and indirect effects were significant, partial mediation was concluded.

## Results

### Study 1: Literature review findings

We reviewed the findings of 23 published studies that utilized graph measurements in aging research, the findings of studies using global and network measures summarized in Table 2. Of the sixteen studies tested global properties, three reported higher clustering coefficient with aging (Sala-Llonch et al., 2014; Xu et al., 2015, Shah et al., 2018), while three found no significant effects (Hugenschmidt et al., 2014; Hou et al., 2019; Yu et al., 2025). Additionally, six papers examined characteristic path length. Three reported higher values with aging (Sala-Llonch et al., 2014; Xu et al., 2015; Mancho-Fora et al., 2020), two reported lower values (Shah et al., 2018; Bagarinao et al., 2019), and one found no significant differences (Hugenschmidt et al., 2014). Age-related alterations in Small-worldness were reported in six studies. Reduced values were reported in three studies (Onoda & Yamaguchi, 2013; Xu et al., 2015; Wang et al., 2024), one reported increased values (Mancho-Fora et al., 2020), and two studies found no significant results (Sala-Llonch et al., 2014; Yu et al., 2025). The remaining studies applied global efficiency as an indicator of global integration. Some reported reduced global efficiency with age (Achard & Bullmore, 2007; Lee et al., 2016; Chong et al., 2019), others found increased values (Chan et al., 2014; Wang et al., 2015; Shah et al., 2018; Bagarinao et al., 2019; Hou et al., 2019), and some reported no significant age-related effects (Cao et al., 2014; Geerligs et al., 2015; Yu et al., 2025). Local efficiency was also used as an indicator of local segregation in seven studies. Two reported increased local efficiency with aging (Shah et al., 2018; Wang et al., 2024), whereas five reported decreased local efficiency (Achard & Bullmore, 2007; Cao et al., 2014; Geerligs et al., 2015; Lee et al., 2016; Chong et al., 2019).

**Table 2:** Review of studies using graph theory in aging.

| **Graph theoretical measures** | **Number of studies** | **Older adults> younger adults** | **Older adults< younger adults** | **No significant differences** |
| --- | --- | --- | --- | --- |
| **Global properties** | 16 |  |  |  |
| Small-worldness | 6 | Mancho-Fora et al., 2020 | Onoda & Yamaguchi, 2013; Xu et al., 2015; Wang et al., 2024 | Sala-Llonch et al., 2014; Yu et al., 2025 |
| Clustering coefficient | 6 | Sala-Llonch et al., 2014; Xu et al., 2015, Shah et al., 2018 | - | Hugenschmidt et al., 2014; Hou et al., 2019; Yu et al., 2025 |
| Characteristic path length | 6 | Sala-Llonch et al., 2014; Xu et al., 2015; Mancho-Fora et al., 2020 | Shah et al., 2018, Bagarinao et al., 2019 | Hugenschmidt et al., 2014 |
| Global efficiency | 11 | Chan et al., 2014; Wang et al., 2015; Shah et al., 2018; Bagarinao et al., 2019; Hou et al., 2019 | Achard & Bullmore, 2007; Lee et al., 2016; Chong et al., 2019 | Cao et al., 2014; Geerligs et al., 2015; Yu et al., 2025 |
| Local efficiency | 7 | Shah et al., 2018; Wang et al., 2024 | Achard & Bullmore, 2007; Cao et al., 2014; Geerligs et al., 2015; Lee et al., 2016; Chong et al., 2019 | - |
| **Network properties** | 12 |  |  |  |
| Modularity | 11 | - | Onoda & Yamaguchi, 2013; Betzel et al., 2014; Chan et al., 2014; Cao et al., 2014; Geerligs et al., 2015; Hou et al., 2019; Wright et al., 2021; Madden et al., 2024; Yao et al., 2025 | Meunier et al., 2009; Baeuchl et al., 2019 |
| Participation coefficient (mean) | 3 | Chan et al., 2014; Chong et al., 2019; Madden et al., 2024 | - | - |
| Participation coefficient (network-wise) | 4 | Meunier et al., 2009; Geerligs et al., 2015; Baeuchl et al., 2019; Chong et al., 2019 | - | - |
| **Nodal properties** | 9 |  |  |  |
| Degree centrality (mean) | 2 | Bagarinao et al., 2019 | - | Baeuchl et al., 2019 |
| Betweenness centrality (mean) | 3 | - | Bagarinao et al., 2019; Wright et al., 2021 | Baeuchl et al., 2019 |

At the network level, twelve studies qualified network metrics, the majority of which examined modularity. Nine studies reported decreased modularity in older adults (Onoda & Yamaguchi, 2013; Betzel et al., 2014; Chan et al., 2014; Cao et al., 2014; Geerligs et al., 2015; Hou et al., 2019; Wright et al., 2021; Madden et al., 2024; Yao et al., 2025), whereas two studies found no significant effects (Meunier et al., 2009; Baeuchl et al., 2019). The participation coefficient was examined in six studies. Three reported increased mean overall participant coefficients in older participants (Chan et al., 2014; Chong et al., 2019; Madden et al., 2024), while others found higher participation coefficient in all cortical networks (Chong et al., 2019), or specific networks, including the SMN and VIS networks (Geerligs et al., 2015), as well as in the DMN network (Baeuchl et al., 2019), and the posterior and superior central modules in older adults (Meunier et al., 2009)

At the nodal level, nine studies examined age-related alterations using graph theoretical analysis. Six, five and two papers calculated degree centrality, betweenness centrality, and nodal efficiency, respectively. Some studies reported increased overall degree centrality with aging (Bagarinao et al., 2019), whereas other found no significant effects (Hugenschmidt et al., 2014). Two studies observed decreased overall BC with aging (Bagarinao et al., 2019; Wright et al., 2021), while one study reported no significant results (Baeuchl et al., 2019). Region-specific findings were also reported, lower degree centrality values in older adults were observed in the bilateral insula, medial prefrontal cortex, putamen, and left caudate, whereas higher degree centrality values were found in the right inferior temporal gyrus and left temporal pole (Cao et al., 2014). Another study reported lower degree centrality and betweenness centrality values in the right anterior cingulate cortex (ACC), as well as lower degree centrality in the left ACC, while betweenness centrality in of the left middle frontal cortex was increased in older adults (Lee et al., 2016). Higher degree centrality values in the right superior parietal gyrus and bilateral precentral gyri were also reported in older participants (Behfar et al., 2020). Nodal efficiency was examined in one study (Achard & Bullmore, 2007), which reported lower nodal efficiency values in the left insula, left thalamus, right hippocampus, and right parahippocampal gyrus, right superior and right inferior frontal gyri, bilateral amygdala and bilateral inferior temporal gyrus in older adults. Only one study examined all three nodal properties (Shah et al., 2018); it reported decreased values in the left middle orbitofrontal gyrus and left inferior temporal gyrus, and increased values in the left inferior occipital cortex, right caudate, right superior temporal gyrus, and right middle temporal pole with aging. Overall, the inconsistencies across studies may be attributed to methodological differences, including variations in parcellation schemes, sparsity thresholds, types of adjacency matrices, sample sizes, and software programs used to compute graph metrics (as reported in Table S1).

Five studies have examined the relationship between the executive functioning and graph measures. Yao et al. (2025) found that decreased modular variability was associated with lower fluid intelligence. Higher participation coefficients were positively associated with better performance on the Stroop task (Baeuchl et al., 2019), whereas a negative relationship was observed between the mean participation coefficient and performance in an attention task (Chong et al., 2019). Two studies reported associations between nodal properties and verbal or associative learning, where the degree centrality of the right posterior cingulate cortex (PCC) was positively correlated with associative learning (Lee et al., 2016); however, degree centrality of the right superior parietal and bilateral precentral gyri showed no association with verbal learning and memory (Behfar et al., 2020). Only one study examined the relationship between global network properties and behavioral performance (Chong et al., 2019), reporting a positive association between local efficiency and attention.

### Study 2: Age-related functional reorganization findings during movie-fMRI

#### Reduced Small-worldness values in older adults

To test our first hypothesis, we examined whether graph theoretical properties differed between older and younger adults. At the global level, older adults exhibited higher clustering coefficient (*t* = –2.98, *p_FDR_* = 0.003, *d* = –0.502) and greater characteristic path length (*t* = –4.36, *p_FDR_* < 0.001, *d* = –0.735), compared to younger individuals (Figure 2A, B). These finding suggest an age-related shift in the balance between functional specialization and integration. Conversely, younger adults showed significantly higher small-worldness values than older participants (*t* = 4.02, *p_FDR_* < 0.001, *d* = 0.674, Figure 2C). However, both age groups demonstrated scores greater than one, confirming the presence of small-world architecture in both groups. These results were further validated across broader sparsity thresholds (i.e.,0.2, 0.25, and 0.3), with the main findings remaining consistent (Figure S1).

**Figure 2.**
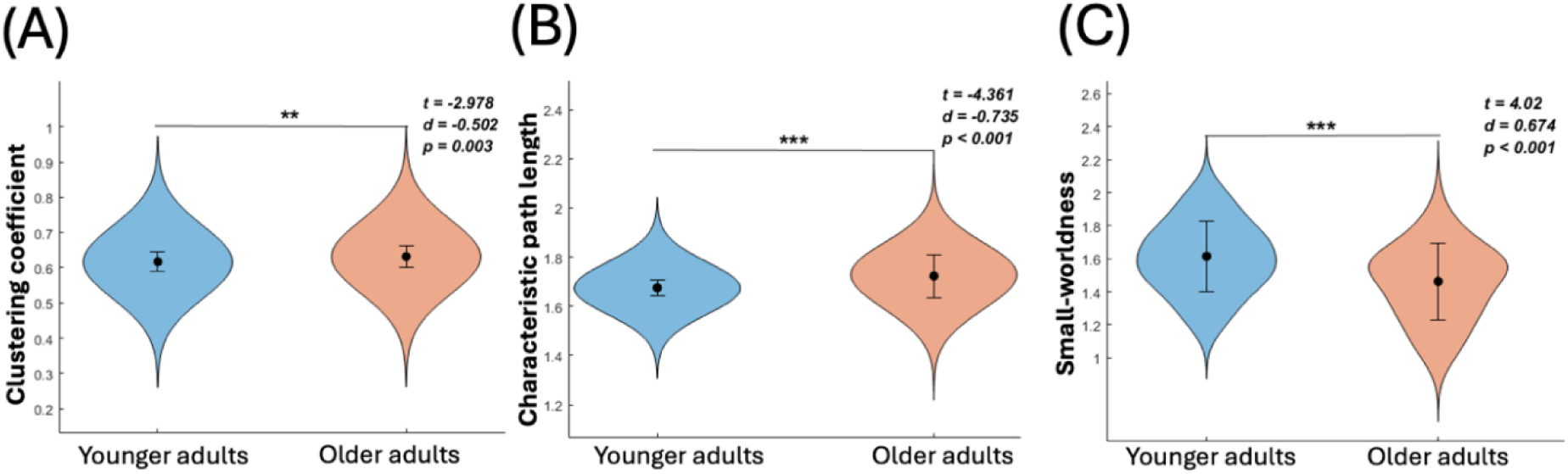
**Age-related differences in global graph properties**. (A) Older adults exhibited significantly higher clustering coefficient, as compared to younger subjects. (B) Older adults demonstrated significantly greater characteristic path length, in comparison with the younger group. (C) Younger adults exhibited significantly higher small-worldness values than older participants. **: *FDR-corrected p < 0.01; ***: FDR-corrected p < 0.001.*

#### Participation coefficient values increased in the SMN, FPN, and DMN networks among older adults

At the network level, the SMN, FPN, and DMN networks exhibited significantly higher participation coefficient values among older adults compared to younger adults (SMN: *t* = –3.21, *p_FDR_* = 0.001, *d* = –0.541; FPN: *t* = –2.77, *p_FDR_* = 0.006, *d* = –0.466; DMN: *t* = –2.59, *p_FDR_* = 0.011, *d* = –0.437), suggesting that these networks showed greater between-network connectivity rather than within-network connectivity in older adults. However, the participation coefficient was higher in the SUB network among younger individuals (SUB; *t* = 3.36, *p_FDR_*= 0.001, *d* = 0.566). In other words, older adults demonstrated greater between-network integration in the sensorimotor, frontoparietal and default mode networks, whereas younger adults showed greater between-network integration of the subcortical network (Figure 3).

**Figure 3.**
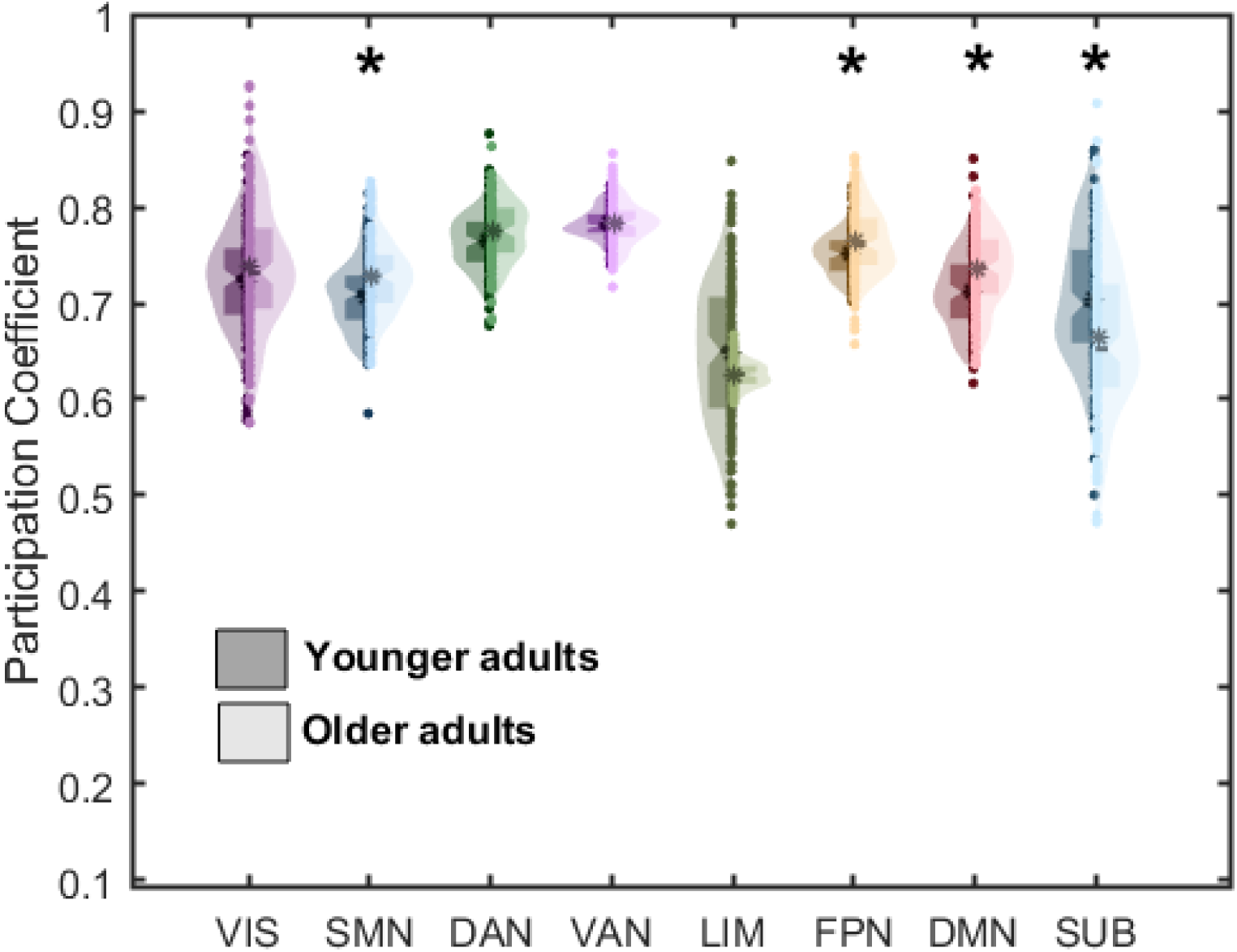
**Age-related differences in network graph properties**. Older adults exhibited significantly higher participation coefficient values in the SMN, FPN and DMN networks, whereas they showed lower participation coefficient values in the SUB network compared to younger adults. VIS= visual network; SMN= somatomotor network; DAN= dorsal attention network; VAN= ventral attention network; LIM= limbic network; FPN=frontoparietal network; DMN= default mode network; SUB= subcortical network; YA= younger adults; OA= older adults. *: *p*-values <0.01, FDR corrected.

#### Lower nodal centrality and efficiency in older adults

At the nodal level, we showed that younger cohort demonstrated significantly higher degree centrality values in several brain regions, including the bilateral hippocampus, bilateral thalamus, right insula, left visual association cortex, right secondary visual areas, right primary auditory cortex, left temporal pole, and left orbitofrontal gyrus. In contrast, older adults showed enhanced degree centrality values in the bilateral dorsolateral prefrontal cortex (DLPFC) and right supramarginal gyrus, compared with the younger adults (all *p*s < 0.001, FDR corrected, Figure 4A). In addition, greater betweenness centrality values in the right hippocampus, bilateral thalamus, and right insula were observed in the younger group. In contrast, lower betweenness centrality values were observed in older adults, specifically in the right primary sensory cortex, left pars orbitalis, and left anterior prefrontal cortex (PFC) (all *p*s < 0.001, FDR corrected, Figure 4B). Moreover, higher nodal efficiency values were observed in the younger cohort in the bilateral hippocampus, bilateral thalamus, bilateral amygdala, bilateral insula, left globus pallidus, left visual association cortex, right secondary visual areas, bilateral primary auditory cortex, left primary sensory cortex, and left inferior temporal gyrus. In contrast, older adults exhibited higher nodal efficiency values in the left DLPFC and right primary sensory cortex (all *ps* < 0.001, FDR corrected, Figure 4C). In summary, five key nodes showed significant group differences across all three properties, including two nodes in the right thalamus, one node in the right hippocampus, and two nodes in the right insula. These regions displayed higher degree centrality, betweenness centrality, and nodal efficiency values in younger adults (as shown in Figure 5A, Table S2).

**Figure 4.**
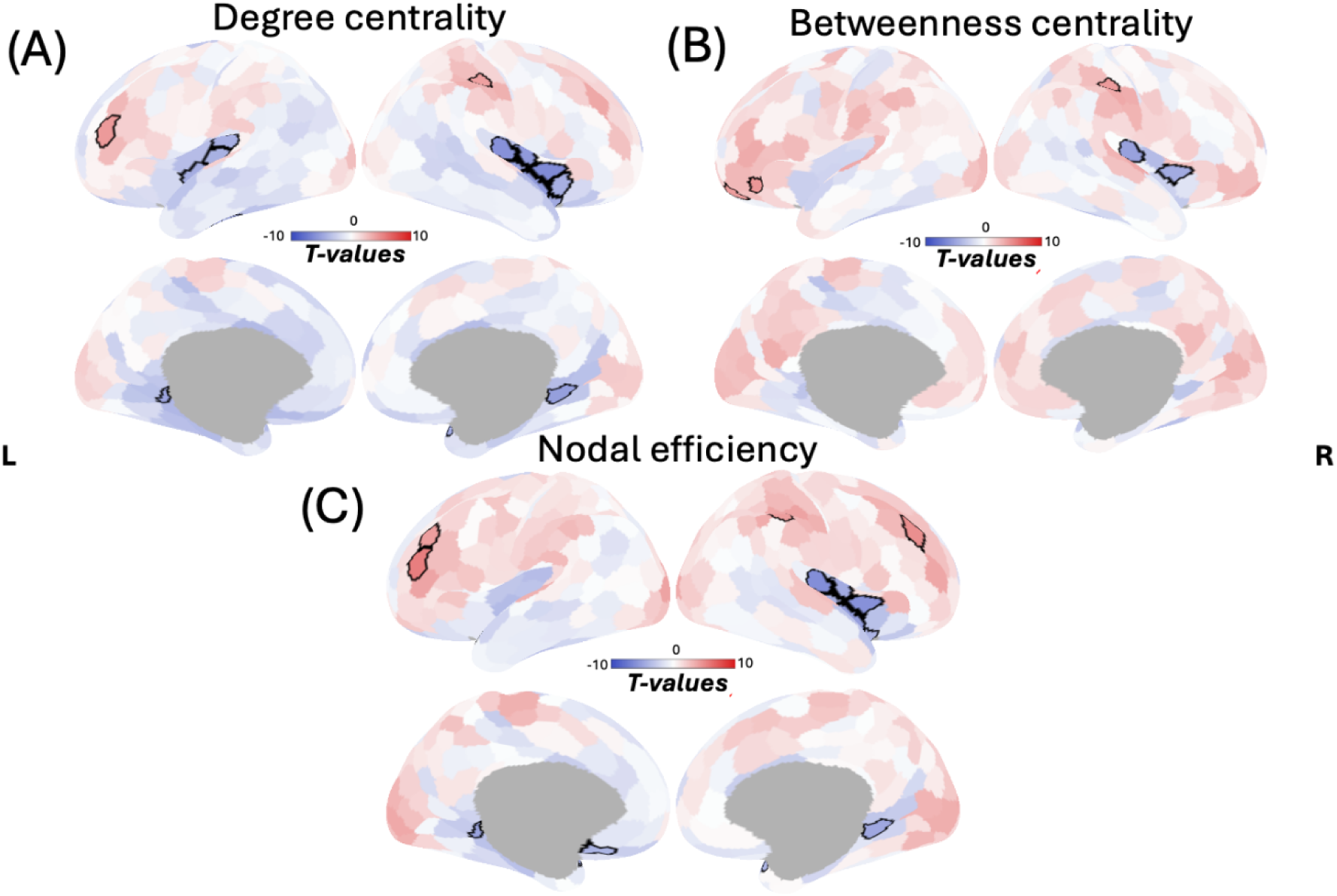
**Age-related differences in nodal graph properties**. Cortical regions in red exhibit higher values of (A) degree centrality, (B) betweenness centrality, and (C) nodal efficiency in older adults, whereas blue-colored regions show higher nodal values in younger adults. Significant contrasts between the two groups are outlined in black. All *p*-values < 0.001, FDR-corrected; L: left hemisphere; R: right hemisphere.

**Figure 5.**
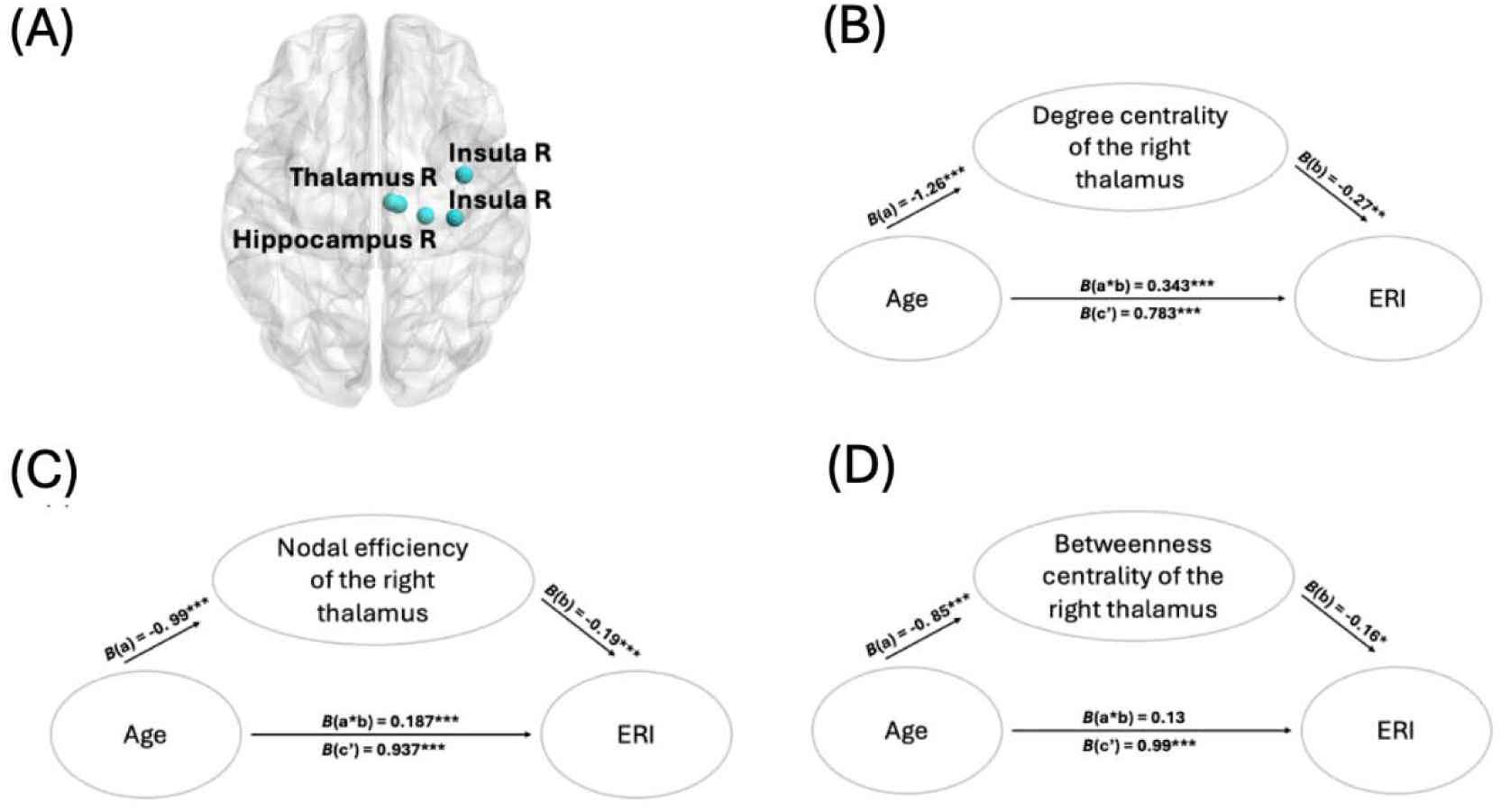
**Key nodes and Partial mediation effects of the degree centrality and nodal efficiency of right thalamus in the relationship of age and ERI**. **(A)** two nodes in the right thalamus, one node in the right hippocampus, and two nodes in the right insula displayed higher degree centrality, betweenness centrality, and nodal efficiency values in younger adults. (B) Lower degree centrality of the right thalamus partially mediates the effect of age on ERI. (C) Lower nodal efficiency of the right thalamus partially mediates the impact of age on ERI. (D) Betweenness centrality of the right thalamus did not mediate the relationship between age and ERI. ERI= emotional resilience index; R: right hemisphere; *: *p* < 0.05, **: *p* < 0.01, ***: *p* < 0.001.

#### Higher nodal properties of right thalamus are negatively associated with ERI

To test our second hypothesis, we further examined whether alterations in graph properties can be associated with emotional and cognitive functioning. The degree centrality, betweenness centrality and nodal efficiency of the right thalamus were significantly associated with ERI (DC: β = –0.27, *t* = –3.04, *p* = 0.003; BC: β = –0.16, *t* = –2.03, *p* = 0.044; NE: β = –0.19, *t* = –2.07, *p* = 0.041, Figure S2A-C), indicating that a higher number of connections and more frequent involvement in efficient pathways via the right thalamus are predictive of lower emotional resilience.

In terms of cognitive reserve, only the degree centrality of the right thalamus showed a marginally significant negative association with CFI (β = –0.19, *t* = –1.92, *p* = 0.057, Figure S2D). Nodal properties of the other key regions, including the right hippocampus and right insula, did not show significant associations with either ERI or CFI. No significant associations were found between global and network graph metrics, including clustering coefficient, characteristic path length, small-worldness, participation coefficient, and the variables of interest, ERI and CFI.

#### Effect of age on ERI are partially mediated by the degree centrality and nodal efficiency of right thalamus

Lastly, given the significant associations between nodal properties of the right thalamus and ERI, we next performed mediation analyses to examine whether these properties mediated the relationship between age and ERI in three separate models. In the first model, age was a significant negative predictor of degree centrality in the right thalamus (path a; *B* = –1.26, SE = 9.38, *p* < 0.001), and degree centrality was negatively associated with ERI (path b; *B* = –0.27, SE = 0.28, *p* = 0.003). The indirect effect of age on ERI through degree centrality was significant (path a × b; *B* = 0.34, 95% CI [0.13, 0.58]), while the direct effect of age on ERI also remained significant (path c′; *B* = 0.78, SE = 3.76, *p* < 0.001, Figure 5B), indicating partial mediation. Thus, reduced degree centrality in the right thalamus partially explained the increased ERI as aging. In the second model, age was negatively associated with nodal efficiency of the right thalamus (path a; *B* = –0.99, SE = 0.02, *p* < 0.001), and lower nodal efficiency was linked to higher ERI (path b; *B* = –0.19, SE = 12.77, *p* < 0.001). The indirect effect of age on ERI via nodal efficiency was significant (path a × b; *B* = 0.19, 95% CI [0.07, 0.36]), and the direct effect of age on ERI also remained significant (path c′; *B* = 0.94, SE = 3.56, *p* < 0.001, Figure 5C), again indicating partial mediation. Therefore, reduced nodal efficiency of the right thalamus partially accounted for the effect of age on ERI. Conversely, betweenness centrality of the right thalamus did not mediate the relationship; the direct effect of age on ERI was significant (path c′; *B* = 0.99, SE = 3.36, *p* < 0.001), but this relationship through the betweenness centrality of right thalamus did not remain significant (path a × b; *B* = 0.13, 95% CI [-0.02, 0.31], Figure 5D).

#### Validation analyses on neutral movie

The changes in graph theoretical metrics in two age groups were replicated while watching the neutral movie. The global measures showed a similar result. There were higher clustering coefficient, higher characteristic path length, and lower small-worldness in older participants. At the network level, all the subnetworks showed increased participation values in older adults. At the regional level, four key nodes, including the bilateral thalamus, left hippocampus and left inferior temporal gyrus indicated higher values in the degree centrality, betweenness centrality, and nodal efficiency in the younger adults. The significant positive association was found between participation coefficient of SMN and ERI, but no mediation effect was observed. Details of results for the neutral movie can be found in the supplementary material.

## Discussion

In the present study, we examined age-related differences in brain network topology during naturalistic movie-watching at global, network, and nodal levels, and tested their associations with emotional and cognitive domains. Aging was associated with systematic alterations in network organization, suggesting a shift in the balance between local specialization and global integration. At the global level, older adults exhibited greater clustering coefficient and longer characteristic path length, indicating greater local segregation but reduced global efficiency. Although both age groups retained small-world organization, reduced small-worldness in older adults suggests a less optimal balance between functional integration and segregation. Reduced small-worldness further suggests diminished global efficiency despite preserved core organizational principles, aligning with prior models of age-related network reconfiguration rather than wholesale network breakdown (Onoda & Yamaguchi, 2013; Deery et al., 2023; Mousley et al., 2025).

At the network level, older individuals showed increased between-module connectivity across sensory (SMN) and higher-order association systems (FPN, DMN), whereas younger adults demonstrated stronger within-module connectivity within the cortical networks and greater between-network integration of subcortical networks. Increased cross-network integration in older adults may reflect age-related network dedifferentiation, characterized by reduced functional specialization and compensatory recruitment across systems. At the nodal level, reduced dominance of hubs was identified in older adults, including the right thalamus, hippocampus and insula. We further found only nodal measures of thalamic areas to show negative association with ERI and CFI and no association with global or network metrics were found. Moreover, the degree centrality and nodal efficiency of the right thalamus partially mediated the relationship between age and ERI, suggesting that decreased centrality of right thalamus partially explains the positive association between age and emotional resilience. Mechanistically, age-related reconfiguration of thalamo-cortical dynamics may preferentially support affective regulation while constraining cognitive flexibility, consistent with socioemotional selectivity and compensatory reorganization frameworks in aging (Carstensen et al., 1999; Cabeza et al., 2002).

Increased characteristic path length in older adults suggests reduced global efficiency. Aligning with the theory of gradual neural pathway degradation (Cabeza et al., 2002), we propose that age-related degradation of long-range white matter pathways perhaps constrains global efficiency, promoting compensatory reconfiguration of shorter-range connections to maintain efficient integration of information (Chakraborty et al., 2024). According to this theory, older adults recruit shorter internal paths to transfer information throughout the brain (Achard & Bullmore, 2007). Prior works has demonstrated life-span related reduction in network efficiency, with increase in early adulthood followed by decline in later life (Achard & Bullmore, 2007; Cao et al., 2014). Variations in the length of neural connections shape transmission dynamics, as long-range projections are metabolically costly and slower to maintain (Chklovskii et al., 2004; Kaiser & Hilgetag, 2006). Thus, increased path length in aging likely reflects reduced long-distance connectivity and greater reliance on multi-step routing, consistent with energy-conserving but less efficient network organization (Achard & Bullmore, 2007).

Consistent with this interpretation, reduced long-range integration may bias networks toward locally dense connectivity, increasing clustering as a compensatory response to diminished global communication. Similarly, increase in clustering have been interpreted as an adaptive mechanism supporting processing efficiency under sensory or structural constraints in older adults, indicating local segregation (Guan et al., 2022). Thus, this pattern reflects a shift toward locally specialized processing at the expense of distributed integration (Mousley et al., 2025). In our results, although older adults exhibited higher local clustering and lower global integration, preservation of small-world structure suggests resilience of core topological principles despite efficiency loss. As denser clustering combined with shorter-range paths with less efficiency in older group, their networks shift toward a more regularized network configuration, resembling patterns observed in Alzheimer’s disease characterized by high segregation and poor global integration (Supekar et al., 2008; Kabbara et al., 2018). This shift may represent a preclinical organizational signature of vulnerability rather than pathology per se.

Aging has been associated with reduced modular segregation and increased between-module connectivity (Geerligs et al., 2014), referred to as dedifferentiation (Park et al., 2004). Reduced modular distinctness in aging may arise from structural atrophy, synaptic loss, and altered functional spatial organization across cortical territories (Cabeza et al., 2002; Krubitzer & Seelke, 2012; Dworetsky et al., 2024). For instance, prior work suggests that the FPN and DMN become less distinct in older adults, functioning more as a unified system (Andrews-Hanna et al., 2007; Geerligs et al., 2014). Such cross-network integration may reflect compensatory recruitment but may also signal reduced between-network specialization critical for flexible cognition. For instance, recent evidence suggests that the FPN network become more integrated in late adulthood to perhaps facilitate the inhibitory control required for emotion regulation in late life (Bätz et al., 2026), although excessive FPN functional dedifferentiation contributes to impaired emotion regulation and poorer mental well-being (Ye et al., 2025). A recent fMRI graph-theoretical study reported decreased participation coefficient of the subcortical module in individuals with high apathy (Zeng et al., 2023). Reduced between-module connectivity between subcortical and cortical systems in aging may therefore reflect altered motivational prioritization, rather than generalized dysfunction, potentially indicating more selective subcortical engagement. Notably, neutral movie did not produce reduced subcortical integration in older adults. This context-dependent effect suggests that subcortical–cortical coupling may be modulated by emotional salience, consistent with adaptive motivational tuning predicted by the SST. Lack of emphasize on subcortical integration in prior rs-fMRI studies may suggest that task demands and affective context, rather than age alone, can impact subcortical integration. Supporting this idea, another study reported that decreased frontal–striatal connectivity was linked to apathy in older adults (Hamada et al., 2021). Altogether, these results may suggest that preserved but more selective subcortical integration may differentiate healthy aging from pathology.

The brain hubs enable long-distance connections to maintain efficient communication (van den Heuvel et al., 2012). It is speculated that nodes distribute their connections more uniformly throughout the network in older adults, which may represent adaptive reweighting of network resources. As a result, network hubs, important for long-distance connectivity, become less prominent with age. Our findings suggest stronger subcortical hub dominance in youth, supporting efficient thalamo-cortical relay and rapid cross-network integration. In healthy aging, there is no dominance of specific hubs, indicating such redistribution may reflect hub dedifferentiation, with less efficient connectivity patterns that weaken long-distance integration and contribute to increased path length. Given the thalamus’ role in balancing cortical integration and segregation (Bell & Shine, 2016; Hwang et al., 2017), altered thalamic centrality likely impacts global network coordination. In our findings, we found that a greater number of connections to the right thalamus, i.e., degree centrality and more efficient connections of the right thalamus with other nodes, i.e., nodal efficiency, were associated with lower emotional resilience. In addition, we found that thalamic centrality and efficiency mediated the association between age and emotional resilience, suggesting that changes in thalamic integration within brain networks may play a critical role in supporting adaptive emotional regulation strategies in older individuals. The thalamus functions as a sensory relay between subcortical and cortical systems, modulates large-scale brain organization (Jones, 1991; Behrens et al., 2003), and its connectivity reflects its involvement in emotional reactivity (Yang et al., 2022). For example, thalamus–amygdala decoupling relates to sensitivity to negative stimuli (Dai et al., 2022), and increased thalamus–ACC connectivity has been observed during encoding of negative material in depression (Lan et al., 2025). Our findings suggest that age-related decreases in thalamic hub dominance, provided by degree centrality and nodal efficiency, may dampen bottom-up affective drive, supporting more stable emotional regulation. Thus, fewer thalamic connections may reflect adaptive processing of salience processing in aging, which also aligns with preferential positive emotional responses in aging (Carstensen et al., 1999), consistent with findings of increased thalamic nodal efficiency in major depression disorder (Ye et al., 2016). Together, our findings suggest that reduced thalamic centrality may prioritize affective salience processing rather than enhance cognitive functioning.

The thalamus also emerged as central for cognitive reserve. Although we did not examine thalamic subregions, posterior thalamus has been implicated in emotion regulation and anterior thalamus in spatial processing (Strigo et al., 2013; Jankowski et al., 2013). Our findings indicated greater right thalamic connectivity that was associated with lower cognitive reserve, consistent with a prior report that increased thalamus–hippocampus coupling relates to poorer cognitive performance in older adults (Goldstone et al., 2018). This pattern may reflect compensatory over-recruitment of subcortical hubs in individuals with reduced cognitive efficiency. While prior work linked right insular centrality to poorer performance in a cognitive task (Lee et al., 2016), our broader PCA-derived cognitive measure may explain the discrepancy. These results indicate that heightened hub centrality does not uniformly provide advantages but may index neural inefficiency under certain conditions.

Several limitations warrant consideration. First, the sample included only healthy older adults, limiting generalizability to clinical populations such as dementia or late-life depression. Second, stimulus variation did not include positive stimuli. Therefore, future work should include positively valanced naturalistic stimuli to examine context-specific network reconfiguration broadly. Third, adjacency matrices were binarized, whereas weighted networks may better capture connection strength (Cole et al., 2010), although this method has been widely used in prior studies (Chan et al., 2014; Cao et al., 2014; Chong et al., 2019; Yao et al., 2025). Future studies should compare differences between the binarized or weighted networks’ approaches. Finally, cross-sectional design limits inference about causality and individual trajectories; longitudinal approaches are needed to follow how these global, network and nodal properties change across time and within-person (Salthouse, 2011).

## Conclusion

The novelty of the present study was on the use of graph theory in combination with the naturalistic movie-fMRI to demonstrate how advancing age influences global, network, and nodal network properties. Our findings provide several novel contributions. In healthy aging, we observed a topological shift toward a more regularized network configuration, i.e., denser local clustering and less efficient long-range connections. We found reduced between-network integration in subcortical network in healthy aging, possibly reflecting altered motivational gating rather than generalized dysfunction, thereby indicating more selective subcortical engagement. Our findings provide novel insight into thalamic importance in global network coordination and topological reconfiguration. Reduced thalamic centrality was associated with better emotional well-being in older adults. These findings suggest that reduced thalamic hub centrality may reflect adaptive recalibration of salience emotional processing, linking network reorganization to improved emotional resilience in aging. These findings highlight the value of a comprehensive, brain-wide approach to functional network topology, offering insights into how aging influences network reconfiguration across multiple levels during naturalistic processing in relation to emotional and cognitive domains.

## Data availability statement

The code and functional connectivity and behavioral data can be found from OSF link: <u>OSF |</u> <u>Brain topology in aging</u>

## Ethics approval statement

This study was conducted in accordance with The Declaration of Helsinki, and ethical approval was granted by the Norwegian Regional Committee for Medical and Health Research Ethics (Midt-REC).

## Patient consent statement

Written informed consent was obtained from all participants.

## Permission to reproduce material from other sources

Figure 1F reproduced without permission from (Farahani FV, Karwowski W and Lighthall NR (2019) Application of Graph Theory for Identifying Connectivity Patterns in Human Brain Networks: A Systematic Review. Front. Neurosci. 13:585. doi: 10.3389/fnins.2019.00585)

## Acknowledgments

We would like to thank radiographers and MR physicists at the 7T MR center at NTNU for their help during this project. We also would like to thank our participants for their time and effort during the experiment. We thank Stian Framvik, Avneesh Jain, Jae Hong, and Karina Tømmerdal for their help during data collection. This project was supported by the Research Council of Norway through its Centers of Excellence scheme, project number 332640 and National Infrastructure grant from the Research Council of Norway (NORBRAIN, project number 245904/350201). We would like to thank Kavli Foundation and hjerneforskningsfondet for their support.

