## supplementary for "Organization of functional brain networks’ architecture during negative movie watching in late adulthood"

These supplementary materials include:

Supplementary Methods

Supplementary Result

Supplementary Fig S1-5

Supplementary Table S1-3

### Supplementary Methods

#### Calculation of graph measures

##### Global properties

Clustering coefficient was calculated based on the following equation:

$c_{i}=\frac{2t_{i}}{k_{i}\left( k_{i}-1 \right)}$	(i)

Where $t_{i}$counts the triangles surrounding node *i,* and $k_{i}$is the number of edges connected to node i. $c_{i}$ denotes the local clustering coefficient. The global clustering coefficient measures the average of triangles across regions obtained based on the following equation:

$C=\frac{1}{N}\sum_{i} c_{i}$	(ii)

Path length was calculated based on the following:

$L=\frac{1}{N}\sum_{i\neq j} L_{ij}$	(iii)

Where N is the total number of nodes, and $L_{ij}$ is the shortest path between nodes *i* and *j*.

Small worldness was calculated as follows:

$S=\frac{C/C_{\text{rand}}}{L/L_{\text{rand}}}$	(iv)

Where *C* is average clustering coefficient, *L* is average characteristic path length, $C_{\text{rand}}$ and $L_{\text{rand}}$are the average clustering coefficient and characteristic path length across 100 random networks. Values above 1 suggest the presence of a small-world network.

##### Network properties

The participation coefficient was calculated as:

$p=1-\sum_{m\in M} \left( \frac{k_{i}\left( m \right)}{k_{i}} \right)^{2}$	(v)

Where *M* indicates a module, $k_{i}\left( m \right)$ is the number of links between node *i* and nodes in module *m*. Values near 1 indicate distributed connections across modules, while values near 0 imply confined connections within a node’s own module.

##### Nodal properties

Degree centrality was calculated as

$D_{i}=k_{i}=\sum_{j} a_{ij}$	(vi)

Where $k_{i}$ equals the total number of connections to node *i*, and the adjacency matrix element $a_{ij}$represents whether node *i* and node *j* are connected or not.

Betweenness centrality was calculated as

$b_{i}=\frac{1}{\left( N-1 \right)\left( N-2 \right)}\sum_{h,j\neq i; h\neq j} \frac{P_{hj}\left( i \right)}{P_{hj}}$	(vii)

Where $\left( N-1 \right)\left( N-2 \right)$ counts the maximum number pairs of nodes not including node *i,* $P_{hj}$ is the total number of shortest paths between node *h* and node *j*, and $P_{hj}\left( i \right)$is the number of those shortest paths passing through node *i*.

Nodal efficiency was calculated as:

$E_{\text{nodal}}\left( i \right)=\frac{1}{N-1}\sum_{j\neq i} \frac{1}{l_{ij}}$	(viii)

Where ${1/l}_{ij}$is the shortest path length between node *i* and node *j.*

#### Supplementary Results

#### Neutral movie

##### Main effect of age in neutral movie

The age-related changes based on graph metrics were replicated while watching the neutral movie. At global level, there were significant differences in clustering coefficient, characteristic path length, and small-worldness with increasing age, while watching neutral. Older adults exhibited a statistically higher clustering coefficient (*t* = –2.73, *p* = 0.007, *d* = –0.46) and higher characteristic path length (*t* = –3.64, *p* < 0.001, *d* = –0.612), compared to younger adults. Moreover, small-worldness is greater than one in both groups, confirming the presence of the balance of functional specialization and integration; however, younger adults showed significantly higher small-worldness values than older adults (*t* = 4.44, *p* < 0.001, *d* = 0.749, Figure S3). These results were further validated across broader sparsity thresholds, with the main findings remaining consistent (Figure S4). These results are similar to the negative movie results.

At the network level, all eight networks exhibited significantly higher participation coefficient values among older adults compared to younger adults. Older adults showed greater integration in the sensory networks (VIS; *t* = –3.44, *p* < 0.001, *d* = -0.781; SMN; *t* = –4.64, *p* < 0.001, *d* = -0.821) , the attention networks(DAN; *t* = –4.61, *p* < 0.001, *d* = -0.94; VAN; *t* = –5.58, *p* < 0.001, *d* = -0.708) , limbic system (LIM; *t* = –4.2, *p* < 0.001, *d* = -0.695), higher-order networks (FPN; *t* = –4.13, *p* < 0.001, *d* = -0.92 ; DMN; *t* = –5.46, *p* <0.001, *d* = -0.977), and subcortical network (SUB; *t* = -5.8, p < 0.001, *d* = -0.58, Figure S5). These results reflect higher integration of all networks while watching the neutral movie in older adults, whereas watching negative movies prohibited the networks losing their variability

At the nodal level, age-related changes in central nodes were also observed while watching the neutral movie. Compared with the older group, the younger cohort demonstrated significantly higher degree centrality in several brain regions, including the bilateral hippocampus, bilateral thalamus, right insula, right primary auditory cortex, right temporal pole, left inferior temporal gyrus, right superior temporal gyrus and right parahippocampus. In contrast, older adults showed enhanced degree centrality in the left DLPFC, left premotor cortex and right anterior PFC. (all *p*s < 0.0001, FDR corrected, Table S3). Additionally, greater betweenness centrality values in bilateral thalamus, left hippocampus, left inferior temporal gyrus, right ventral anterior cingulate cortex, were observed in the younger group. In contrast, lower betweenness centrality was observed in older adults, specifically in the left Broca and right anterior PFC (all *p*s < 0.0001, FDR corrected, Table S3).

Moreover, higher nodal efficiency was observed in the younger cohort in the bilateral hippocampus, bilateral thalamus, right amygdala, bilateral insula, right parahippocampus, left visual association cortex, bilateral temporal pole, right primary auditory cortex, left inferior temporal gyrus, left orbitofrontal cortex, and bilateral superior temporal gyri. Older adults exhibited no significant increased nodal efficiency. (all *p*s < 0.0001, FDR corrected, Table S3). To summarize, four key nodes indicated significant differences in all three properties, including the bilateral thalamus, left hippocampus and left inferior temporal gyrus. These regions displayed higher degree centrality, betweenness centrality and nodal efficiency in younger adults.

##### Brain-behavior relationship during neutral movie

To examine the association of graph metrics with behavioral measures while watching the neutral movie, participation coefficient of SMN showed significant positive relationship with ERI (β = 0.2, *t* = 2.32, *p* = 0.022). Moreover, small-worldness has a marginal negative association with ERI (β = -0.16, *t* = -1.91, *p* = 0.058), whereas marginal positive association was observed between betweenness centrality of left inferior temporal gyrus and ERI (β = 0.15, *t* = 1.95, *p* = 0.054). In terms of cognitive reserve, only the participation coefficient of VIS and DAN showed a marginally significant negative association (β = –0.17, *t* = –1.92, *p* = 0.056; β = –0.17, *t* = –1.94, *p* = 0.055). No mediation analyses were significant.

Table S1: Review of previous papers using graph theory methods to investigate age-related alternations.

| # | **First Author, Year** | **Field strength** | **fMRI** | **Atlas** | **Software for graph analysis** | **Contrast** | **Network**  **properties** | **Threshold range** | **Type of graph** |
| --- | --- | --- | --- | --- | --- | --- | --- | --- | --- |
| **1** | ***Achard et al., 2007*** | - | Rs-fMRI | AAL (90 Nodes) | Brainwaver | OA(N=13) vs. YA (N=17) | GE, LE, NE | 0.05: 0.1:0.35 | B |
| **2** | ***Meunier et al., 2009*** | 3T | Rs-fMRI | AAL (90 Nodes) | n.a. | OA(N=13) vs. YA (N=17) | Q, PC | Absolute 0.05 | B |
| **3** | ***Onoda et al., 2013*** | 1.5T | Rs-fMRI | AAL (90 Nodes) | BCT | Age effect  (N=193) | Q, Sigma | n.a. | B |
| **4** | ***Sala-Llonch et al., 2014*** | 3T | Rs-fMRI | AAL (90 Nodes) | BCT | Age effect  (N=98) | Sigma, GE, Lp, CP | Absolute 0.15 | B |
| **5** | ***Betzel., 2014*** | 3T | Rs-fMRI | 114 nodes | BCT | Age effect  (N=126) | Q | Not thresholded | W |
| **6** | ***Chan et al., 2014*** | 3T | Rs-fMRI | 441 nodes | BCT | Age effect  (N=210) | Q, GE, PC | 0.3:0.01:0.1 | W |
| **7** | ***Cao et al., 2014*** | 3T | Rs-fMRI | LYeo131 (1024 nodes) | n.a. | Age effect  (N=126) | M, GE, LE, DC | 0.05 : 0.2 | W, B |
| **8** | ***Hugenschmidt et al., 2014*** | 1.5 T | Rs-fMRI | AAL (90 Nodes) | n.a. | OA(N=48) vs. YA (N=24) | Lp, Cp, DC | 0.2:0.05:0.3 | B |
| **9** | ***Geerligs et al., 2015*** | 3T | Rs-fMRI | Power atlas (264 nodes) | BCT | OA(N=40) vs. YA (N=40) | Q, GE, LE, PC | Absolute 0.27 | B |
| **10** | ***Xu et al., 2015*** | - | Rs-fMRI | AAL (90 Nodes) | n.a. | OA(N=11) vs. YA (N=195) | Sigma, Lp, Cp | 0.05:0.05:0.3 | B |
| **11** | ***Lee et al., 2016*** | 3T | Rs-fMRI | n.a. | n.a. | Age effect  (N=155) | GE, LE, DC, BC | Absolute 0.21 | B |
| **12** | ***Shah et al., 2018*** | 3T | Rs-fMRI | AAL (90 Nodes) | Gretna | Age effect  (N=458) | DC, BC, NE, GE, LE, Lp, CP | Absolute 0.2 | W |
| **13** | ***Hou et al., 2019*** | 3T | Rs-fMRI | 114 nodes | BCT | Age effect  (N=170) | GE, Cp, Q | n.a. | B |
| **14** | ***Baeuchl et al., 2019*** | 3T | Rs-fMRI | 200 nodes | BCT | OA(N=37) vs. YA (N=41) | Q, PC, BC | 0.3:0.45 | B |
| **15** | ***Chong et al., 2019*** | 3T | Rs-fMRI | 114 nodes | BCT | OA(N=72) vs. YA (N=57) | GE, LE, PC | 0.1:0.01:  0.3 | W |
| **16** | ***Bagarinao et al., 2019*** | 3T | Rs-fMRI | 499 nodes | Gretna | Age effect  (N=129) | GE, Lp, BC | Absolute 0.2 | B |
| **17** | ***Behfar et al., 2020*** | 3T | Rs-fMRI | Brainnetome atlas (246 nodes) | Conn toolbox | OA(N=15) vs. YA (N=15) | S | 0.05:0.01:0.5 | B |
| **18** | ***Mancho-Fora al., 2020*** | 3T | Rs-fMRI | AAL (90 Nodes) | n.a. | Age effect  (N=114) | Sigma, Lp, Cp | n.a. | B |
| **19** | ***Wright et al., 2021*** | 3T | Rs-fMRI | 16 nodes | BCT | OA(N=70) vs. YA (N=75) | GE, LE, Cp, Q, BC | Absolute 0.05 | B |
| **20** | ***Madden et al., 2024*** | 3T | Rs-fMRI | Brainnetome atlas (228 nodes) | BCT | Age effect  (N=68) | Q, PC | n.a. | B |
| **21** | ***Wang et al., 2024*** | - | Rs-fMRI | Power atlas (226 nodes) | Gretna | Age effect  (N=998) | Sigma, GE, LE, Lp, Cp | 0.05:0.01:0.4 | B |
| **22** | ***Yu et al., 2025*** | 3T | Rs-fMRI | Schaefer 200 atlas + 19 subcortical areas | BCT | OA(N=36) vs. YA (N=41) | Sigma, GE, Cp | Absolute 0.2 | B |
| **23** | ***Yao et al., 2025*** | - | Rs-fMRI | Power atlas (264 nodes) | n.a. | Age effect  (N=615) | Q | 0.1:0.05:0.15 | W |

Note: Rs-fMRI: resting-state functional magnetic resonance imaging; BCT: Brain connectivity toolbox; OA: Older adults; YA: Younger adults; n.a.: not available; Sigma: Small-worldness; Lp: Characteristic path length; Cp: Clustering coefficient; GE: Global efficiency; LE: Local efficiency; Q: Modularity; PC: Participation coefficient; DC: Degree centrality; BC: Betweenness centrality; NE: Nodal efficiency; B: Binary; W: Weighted.

**Table S2**: Differences of younger and older adults in nodal graph properties while watching the negative movie.

| Brain regions | Degree centrality | |  | Betweenness centrality |  | Nodal efficiency |
| --- | --- | --- | --- | --- | --- | --- |
|  | *t*-value | |  | *t*-value | *t*-value | |
| **Thalamus R** | 9.52*** | |  | 5.75*** | 8.45*** | |
| Thalamus L | 7.33*** | |  |  | 6.91*** | |
| **Hippocampus R** | 5.83*** | |  | 4.9*** | 4.73*** | |
| Hippocampus L | 6.39*** | |  |  | 5.8*** | |
| Amygdala R |  |  |  |  | 5.2*** | |
| Amygdala L |  |  |  |  | 4.47*** | |
| Globus pallidus L |  |  |  |  | 4.31*** | |
| Visual association L | 5.02*** | |  |  | 4.91*** | |
| Secondary visual R | 4.81*** | |  |  | 5.28*** | |
| Primary auditory R | 4.26*** | |  |  | 5.17*** | |
| Primary auditory L |  |  |  |  | 4.26*** | |
| Primary sensory R |  |  |  | -4.26*** | -4.09*** | |
| Primary sensory L |  |  |  |  | 4.35*** | |
| **Insula R** | 6.21*** | |  | 5.15*** | 6.4*** | |
| Insula L |  |  |  |  | 4.47*** | |
| Temporal pole L | 4.38*** | |  |  |  | |
| Inferior temporal L |  |  |  |  | 5.51*** | |
| Orbitofrontal L | 4.13*** | |  |  |  | |
| Anterior PFC R |  |  |  | -4.23*** |  | |
| DLPFC R | -4.38*** | |  |  |  | |
| DLPFC L | -5.56*** | |  |  | -4.36*** | |
| Supramarginal gyrus R | -4.25*** | |  |  |  | |
| Pars orbitalis L |  |  |  | -4.48*** |  | |

Note: *** Indicates regions with significant *p*-values in all nodal properties (degree centrality, betweenness centrality and nodal efficiency). PFC: prefrontal cortex; DLPFC: dorsolateral prefrontal cortex; L: left hemisphere; R: right hemisphere.

**Table S3:** Differences of younger and older adults in nodal graph properties while watching the neutral movie.

| Brain regions | Degree centrality | | Betweenness centrality |  | Nodal efficiency | |
| --- | --- | --- | --- | --- | --- | --- |
|  | *t*-value | | *t*-value |  | | *t*-value |
| **Thalamus R** | 7.67*** | | 3.99*** |  | | 7.69*** |
| **Thalamus L** | 6.45*** | | 6.67*** |  | | 6.55*** |
| Hippocampus R | 5.47*** | |  |  | | 5.46*** |
| **Hippocampus L** | 8.22*** | | 7.18*** |  | | 7.43** |
| Parahippocampus R | 4.03*** | |  |  | | 4.31** |
| Amygdala R |  |  |  |  | | 4.31** |
| Visual association L |  |  |  |  | | 4.15** |
| Premotor L | -4.63*** | |  |  | |  |
| Primary auditory R | 4.35*** | |  |  | | 5.11*** |
| Insula R | 6.1*** | |  |  | | 6.18*** |
| Insula L |  |  |  |  | | 4.47*** |
| Temporal pole R | 3.9*** | |  |  | | 4.42*** |
| Temporal pole L |  |  |  |  | | 4.28*** |
| **Inferior temporal L** | 5.01*** | | 5.36*** |  | | 5.42*** |
| Superior temporal R | 4.44*** | |  |  | | 5.25*** |
| Superior temporal L |  |  |  |  | | 4.29*** |
| Orbitofrontal L |  |  |  |  | | 4.48*** |
| Anterior PFC R | -4.09*** | | -3.93*** |  | |  |
| DLPFC L | -4.02*** | |  |  | |  |
| L Broca |  |  | -4.08*** |  | |  |
| vACC R |  |  | 3.96*** |  | |  |

Note: *** Indicates regions with significant *p*-values in all nodal properties (degree centrality, betweenness centrality and nodal efficiency. PFC: prefrontal cortex; DLPFC: dorsolateral prefrontal cortex; vACC: ventral anterior cingulate cortex; L: left hemisphere; R: right hemisphere.

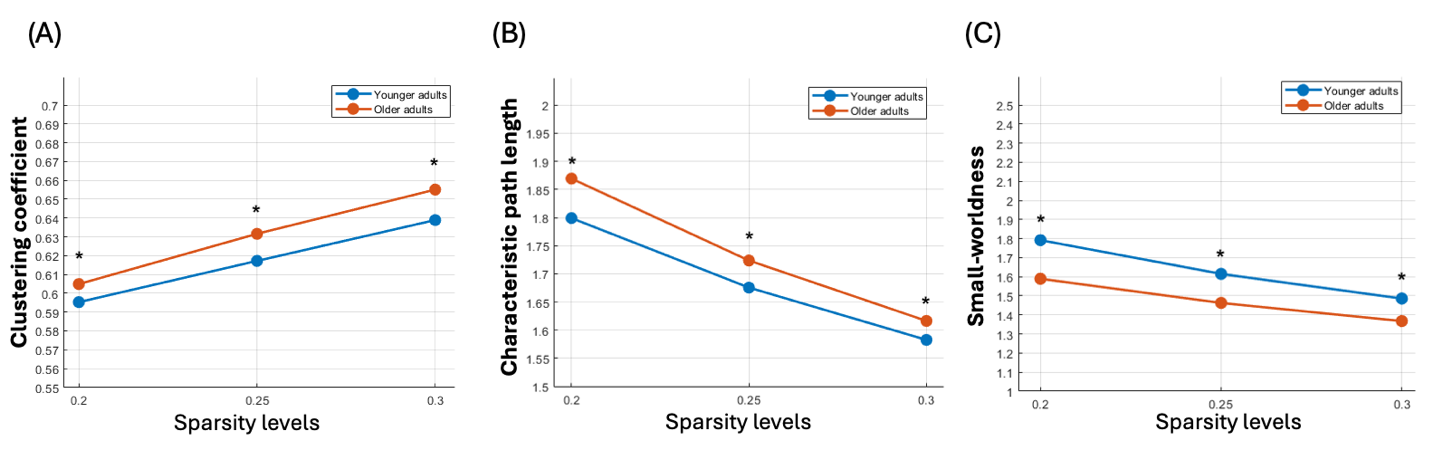

**Figure S1.** Global properties exhibited significant differences between younger and older adults during negative movie watching, on broader sparsity thresholds (0.2, 0.25, 0.3). A) Clustering coefficient values, (B) characteristic path length values, and (C) small-worldness values. Blue indicates younger adults and green indicates older adults. All *p*-values<0.05

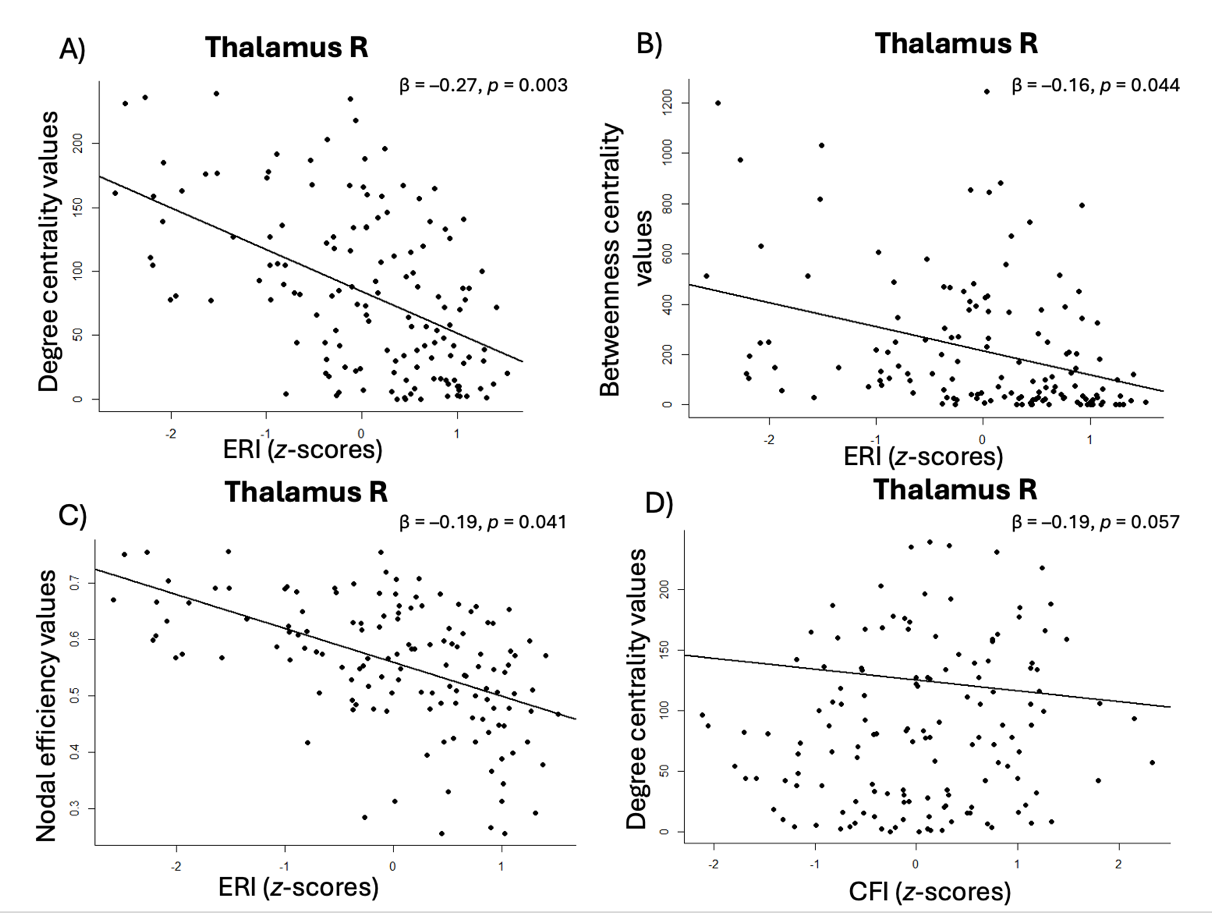

**Figure S2.** Scatterplots of the multiple linear regression analyses between nodal properties of the thalamus ERI. (A) Degree centrality of the right thalamus is negatively associated with ERI, (B) betweenness centrality of the right thalamus is negatively associated with ERI, (C) nodal efficiency of the right thalamus is negatively associated with ERI. (D) Degree centrality of right thalamus is negatively associated with CFI. All *p*-values < 0.001 FDR-corrected; R: right hemisphere..

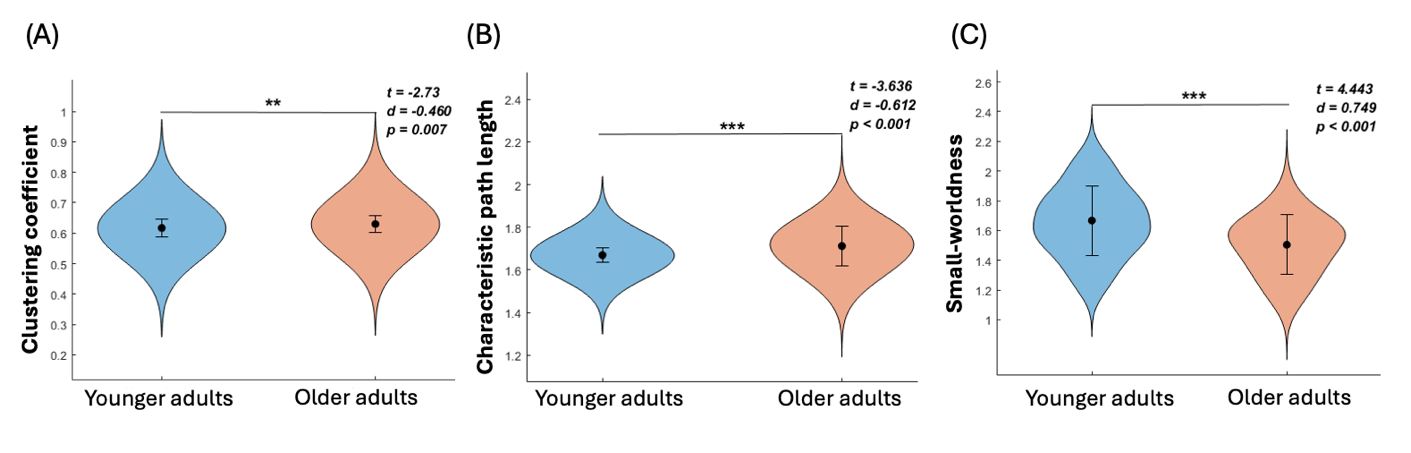

**Figure S3.** Age-related differences in global graph properties, comparing older adults with younger adults during watching neutral movie. (A) Older adults exhibited significantly higher clustering coefficient, as compared to younger subjects. (B) Older adults demonstrated significantly greater characteristic path length, in comparison with the younger group. (C) Younger adults exhibited significantly higher small-worldness values than older participants. **: *p < 0.01; ***: p < 0.001.*

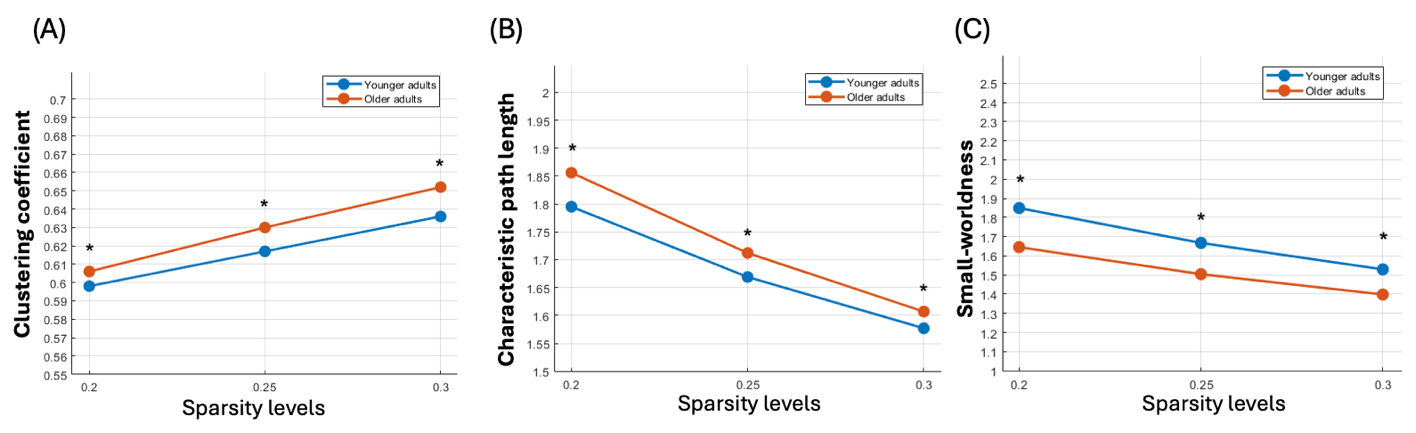

**Figure S4:** Global properties exhibited significant differences between younger and older adults during neutral movie watching, on broader sparsity thresholds (0.2, 0.25, 0.3). A) Clustering coefficient values, (B) characteristic path length values, and (C) small-worldness values. Blue indicates younger adults and green indicates older adults. All *p*-values<0.05

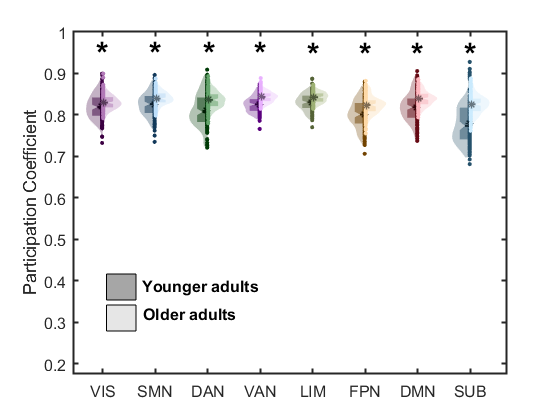

**Figure S5.** Age-related differences in the modular graph property, comparing older adults with younger adults while watching the neutral movie. Older adults exhibited higher participation coefficient values in all the networks, as compared to younger adults. VIS= visual network; SMN= somatomotor network; DAN= dorsal attention network; VAN= ventral attention network; LIM= limbic network; FPN=frontoparietal network; DMN= default mode network; SUB= subcortical network; YA= younger adults; OA= older adults. *: *p*-values <0.01, FDR corrected.
